# The Restoration of Social Defects in Schizophrenic Mice by Plant Exposure

**DOI:** 10.1101/2024.12.23.630193

**Authors:** Mingxuan Li, Mingda Li, Kai Zhou, Xiangyu Zhang, Shanshan Liu, Yuyu Wu, Xinjian Li

## Abstract

Schizophrenia is a serious psychotic disorder caused by both individuals’ genetic background and their living environment. However, how the cognitive and negative symptoms can be treated is still a challenge since most anti-psychotic drugs are only effective for positive symptoms. Previous epidemic studies have demonstrated that plant exposure could decrease the risk of schizophrenia and shorten the length of psychiatric hospital admissions for patients. However, it is still unknown whether plant exposure could improve cognition-related defects in schizophrenia, with no related animal studies. In the present study, we first induced a schizophrenia mice model by giving mice long-term (2 weeks) injections of the antagonist of the NMDA receptor: MK801. We then raised the animals in environments containing plants (4 weeks) including Epipremnum *aureum* and rosemary, and tested their locomotive, anxious and social behaviors. We found that plant exposure did not change the locomotion behaviors of wild-type animals, but significantly reduced the schizophrenia-related social and anxious behaviors in the schizophrenic animals. In addition, we tested the expression of c-Fos in animals exposed to plants after social behavioral testing and found that the deficits in c-Fos expression in both the hippocampus and prefrontal cortex were partially rescued after plant exposure. These results indicate plant exposure will be a new tool to improve the clinical deficits of schizophrenia.

## Introduction

Schizophrenia, a severe mental disorder, impacts 0.5-1% of the global population, significantly disrupting the lives of individuals and potentially leading to social disaters^1,2.^ Through advancements in genetic and molecular research, schizophrenia has been associated with certain genetic mutations and environmental influences, including those that affect individuals before birth. Despite the diverse origin of schizophrenia, individuals with the disease share several common symptoms that can be grouped into 3 clusters: positive, negative, and cognitive dysfunction^1,2^. Positive symptoms typically relate to symptoms associated with a “break from reality”, such as delusions, hallucinations, incoherent speech, and erratic behavior. Negative symptoms are typically identified by amotivational syndrome, which includes lack of drive, affective flattening, social avoidance, anhedonia, and diminished energy. The cognitive symptoms of schizophrenia manifest as a broad set of cognitive dysfunctions^1^. Though many antipsychotic medications have been developed in the past three decades to manage the clinical manifestations of schizophrenia, most are only effective in alleviating positive symptoms. Although cognitive symptoms also significantly impair patients’ daily lives, few treatment methods have been clinically available until now.

Under natural conditions, humans and animals live together with many different kinds of plants, forming symbiotic relationships. Plants perform photosynthesis, producing oxygen and synthesizing glucose. Additionally, plants also emit different smells and visual stimuli, which can improve the sensory, emotional, and social behaviors of humans and animals. In return, animals provide organic manure and help to shape plant morphology. So, plant exposure is beneficial for both animals and plants. Previous studies have indicated that abnormal environments during pregnancy and childhood development could be one of the key factors in the development of schizophrenia. People residing in urban cities are at a higher risk of developing schizophrenia compared to those living in rural regions.^3^. A contributing factor to this phenomenon may be that urban areas contain fewer plants than rural areas. Several previous studies have indicated that green space exposure decreased the risk of schizophrenia in a dose-dependent manner to some extent^4 5-7^ and can shorten the length of psychiatric hospital admissions ^8^. So, we hypothesize that enough plant exposure may be able to rescue the cognitive symptoms of schizophrenia. By raising schizophrenic mice in environments containing plants, we found schizophrenia-related social and anxiety deficits are rescued. In addition, the neural activity related c-Fos expression was partly rescued in both the prefrontal cortex and hippocampus of schizophrenic mice after plant exposure. These findings provide a possible novel approach for treating the cognitive and negative symptoms associated with schizophrenia.

## Methodology

### Subjects

Adult male C57BL/6 mice (8-10 weeks,20-25g) were used for all experiments. The mice were housed in large plastic cages (0.7m x0.7m x0.6m), with 8-9 mice per cage. The room temperature was kept at 25 ± 1 °C, and lighting was set to a 12 h light/dark cycle. The humidity of the room is around 40 ± 5%. The Animal Use and Care Committee of Zhejiang University approved all experimental procedures.

### Animal modeling

The schizophrenic mice were modeled based on previous studies^9,10^, MK-801 (0.05mg/ml) was injected (I.p) to C57BL/6 mice at the dosage of 0.5mg/kg twice per day. The control mice were injected with saline twice per day at the same volume. The injections lasted for two weeks.

### Groups

The animals were separated into 4 groups (**Fig. 1D**): wild type (WT) raised without plants (Sch-/ Plant-, n=9), WT with plants (Sch- / Plant+, n=8), schizophrenia without plants (Sch+ / Plant-, n=8), schizophrenia with plants (Sch+ / Plant+, n=8).

**Figure 1.**
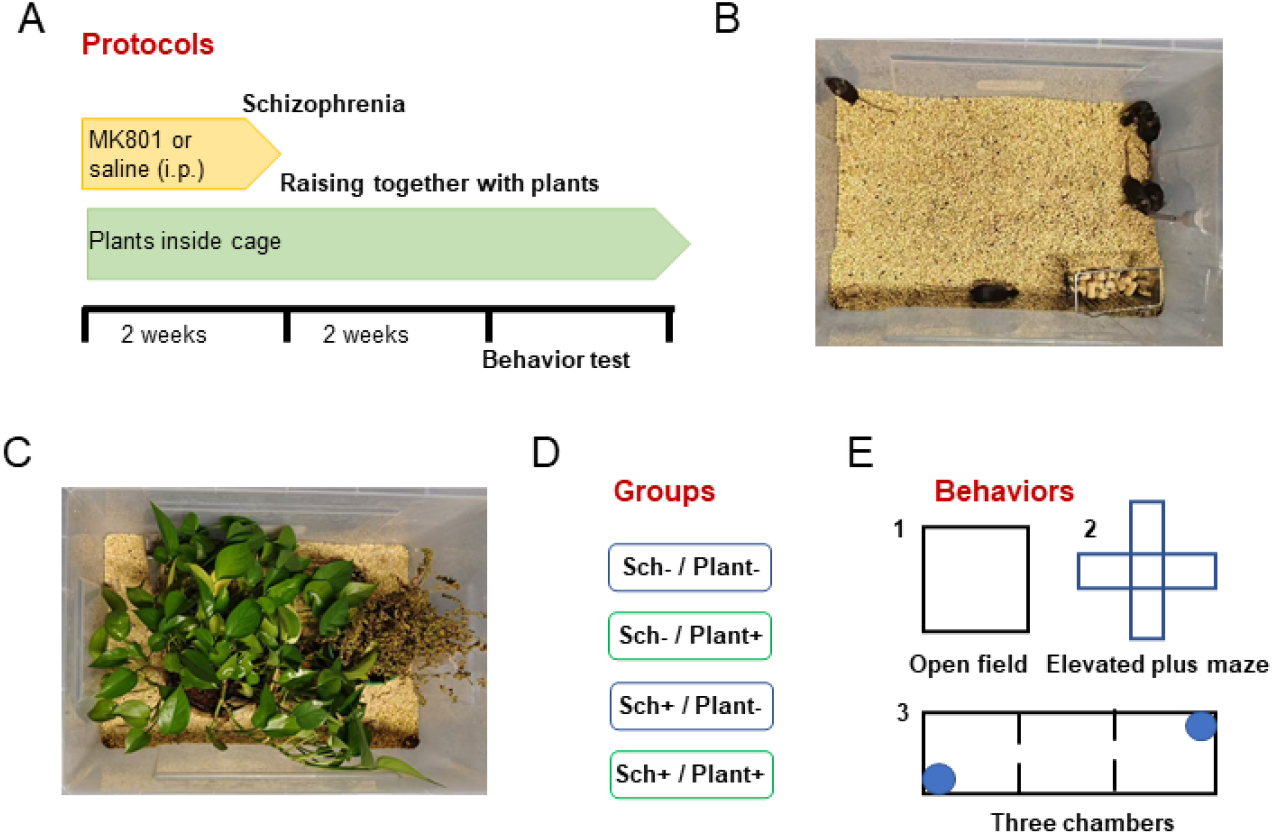
Plant exposure protocol. (A) The protocol for plant exposure. Upper panel: to model schizophrenia, MK801 (i.p., 0.5 mg/kg) or saline was administered to WT (C57B/L6) mice via intraperitoneal injection (i.p.) over two weeks. Middle, the timeline for raising the animals together with plants (4 weeks). Bottom, the timeline for modeling schizophrenic animals, plant exposure and behavioral testing. (B) An example imaging showing that the animals were housed in a 0.7m x 0.7m plastic box without plants. (C) An example imaging showing that the animals were housed in a 0.7m x 0.7m plastic box containing plants. (D) The different groups of animals. Sch- / Plant-, wild type (WT) raised without plants; Sch- / Plant+, WT with plants; Sch+ / Plant-, schizophrenia without plants; Sch+ / Plant+, schizophrenia with plants. (E) The behavioral tests conducted in sequence and separated in different days, including 1. the open field, 2. the elevated plus maze, and 3. the three-chamber social test.

### Plant exposure

In this study, we built environments containing plants to address whether plant exposure can alleviate social-related symptoms in schizophrenia. Two kinds of plants including *Epipremnum aureum* and rosemary were chosen. *Epipremnum aureum* is one of the most popular home pot plants, with strong leaves and great survivability. Rosemary (*Salvia rosmarinus*) is a small woody shrub that includes benefits such as antimicrobial and antioxidant properties and is used as an alternative method to treat mental fatigue, etc. In the present study, we put both rosemary and *Epipremnum aureum* inside the plastic cages. Because of the shape of the plants, the animals could run between and hide in the plants and even eat some parts of the plants. In addition, we observed that the mice exposed to plants became wild and had a stronger willingness to interact with the environment and other animals. By raising the mice in environments containing plants for 4 weeks, we aimed to dissect whether plant exposure could improve social-related defects in schizophrenic mice.

### Behavior test

In this present study, we evaluated emotion-related behaviors using the open field and elevated plus maze test, and assessed social behaviors with the three-chamber social test. The animals were first introduced to the experimental room, where they spent 10 minutes daily for 3 consecutive days to acclimate to the environment and become familiar with the handling of the experimenter. For the open field test, each mouse was placed in the center of the field at the start of each trial and allowed 8 minutes of free exploration. In the elevated plus maze, each mouse was initially placed on one of the open arms and allowed 8 minutes of free exploration. The three-chamber social test was carried out in three phases: habituation, social ability testing, and social novelty testing, each phase lasting 10 minutes. In the habituation phase, no animals were placed inside the pen cups, and the tested mouse was allowed to explore the three-chamber freely. Then, in the second phase (social ability), a novel conspecific Stranger 1, who is the same sex and strain as the subject, is placed in cup 1. In the third phase (social novelty), a second novel conspecific Stranger 2, who is also the same sex and strain, is placed under cup 2. Stranger 1 remains in cup 1. Between each trial or phase of every test, a 75% alcohol solution was sprayed to remove the traces of scent left by previous test subjects. All behavioral video was recorded at 60 frames per second with a high-definition camera (1080p, Ordro, version D395).

### Histology

Immunohistochemistry was performed as described in a previous study^11^. To quantify the c-Fos positive cells, we sampled sections every 80-μm from the whole brain of mice from each of the 4 groups. Brain regions were outlined and compared according to the reference atlas (the Allen Mouse Brain Atlas). And c-Fos positive cells were quantified using ImageJ.

### Statistical Analyses

The bar graphs are presented as the mean plus the standard error of the mean (SEM). Statistical comparisons between groups were performed using either one-way or two-way ANOVAs (analysis of variance), followed by a rank-sum test for multiple comparisons.

## Results

### Plant exposure does not change the locomotive behaviors in either wild-type or schizophrenic mice

To evaluate whether plant exposure could improve behavioral deficits in animals with mental disorders, we first induced a schizophrenia mice model by giving the mice long-term (2 weeks) injections of the antagonist of the NMDA receptor: MK801(i.p.,0.5mg/kg, 2 weeks). We then raised the mice in environments containing two species of plants, *Epipremnum aureum* and rosemary, for at least two weeks (**Fig. 1A-C**). As they were raised, the mice could actively interact with plants, including hiding inside, running between, and even eating some parts of the plants (data not shown). The animals were separated into 4 groups (**Fig. 1D**): wild type (WT) raised without plants (Sch- / Plant-, n=9), WT with plants (Sch- / Plant+, n=8), schizophrenia without plants (Sch+ / Plant-, n=8), schizophrenia with plants (Sch+ / Plant+, n=8). Then, we assessed the social, anxious and locomotive behaviors (Fig. 1E).

MK801 is an NMDA receptor antagonist and may greatly change the locomotive behavior of the mice after long-term administration. Therefore, we evaluated their mobility by analyzing their movements in the open filed test (**Fig. 2A**). The mice’s running trace was recorded and overlain onto the open field. As shown in Figure 2A, the mice ran across almost all of the positions in the open field. Then we analyzed the averaged running distance (**Fig. 2B**) and speed (**Fig. 2C**) over 8 minutes of testing for each of the 4 groups of mice. We found that plant exposure did not change the locomotive behaviors in either wild-type or schizophrenic mice in the open field (**Fig. 2B-2C**).

**Figure 2.**
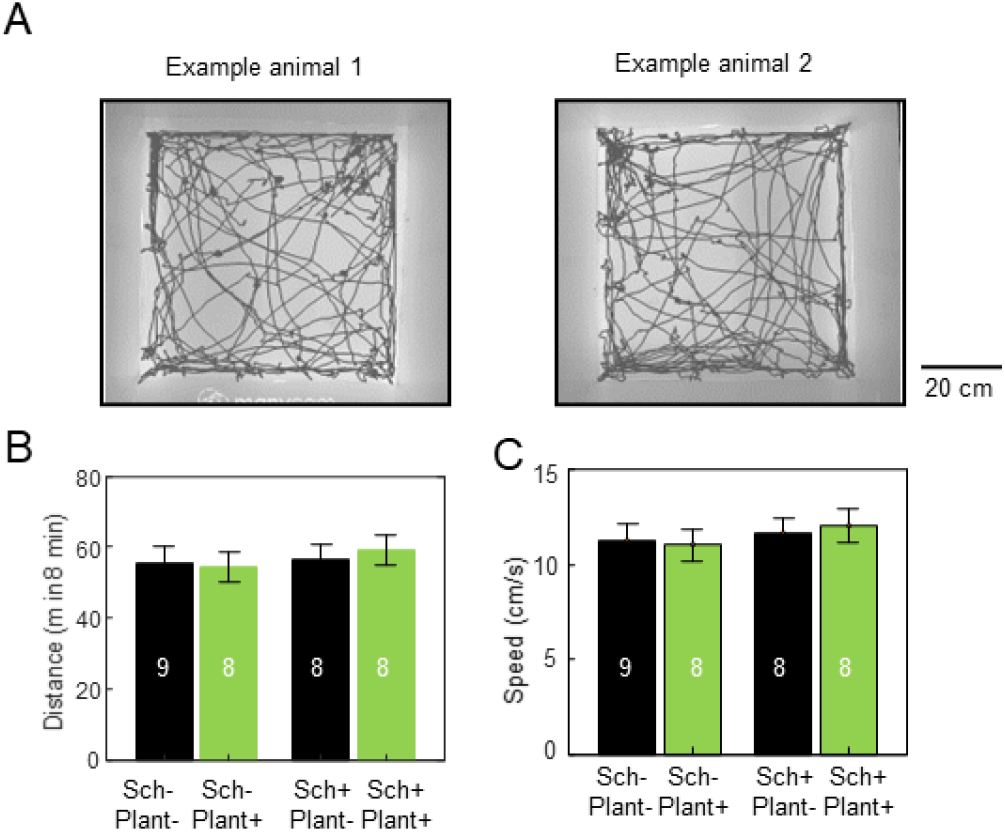
Plant exposure does not change the locomotive behaviors in either WT or schizophrenic mice. (A) The 8-minute walking trace of two representative mice in the open field, with movement trajectories covering most areas of the open field. (B) The average running distance of the four groups of animals over 8 minutes. (C) The average running speed of the four groups of animals over 8 minutes.

As we know, most mental diseases including schizophrenia will induce anxiety^12-14^. To test whether plant exposure can rescue anxiety deficits caused by MK801-induced schizophrenia, we first analyzed the time of each mouse spent running in the center and corner segments of the open field (**Fig. 3A and 3B**). We observed that the schizophrenic mice spent less time running in the center of the field in comparison to the WT mice (**Fig. 3C**). Importantly, plant exposure significantly increased the running time in the center compared with the animals raised without plants (**Fig. 3C**). However, the increase in center running time was not shown in the WT animals with plant exposure (**Fig. 3C**). So, plant exposure significantly decreased the schizophrenia-related anxious behavior of mice in the open field test, and had little effect on WT mice’s anxiety.

**Figure 3.**
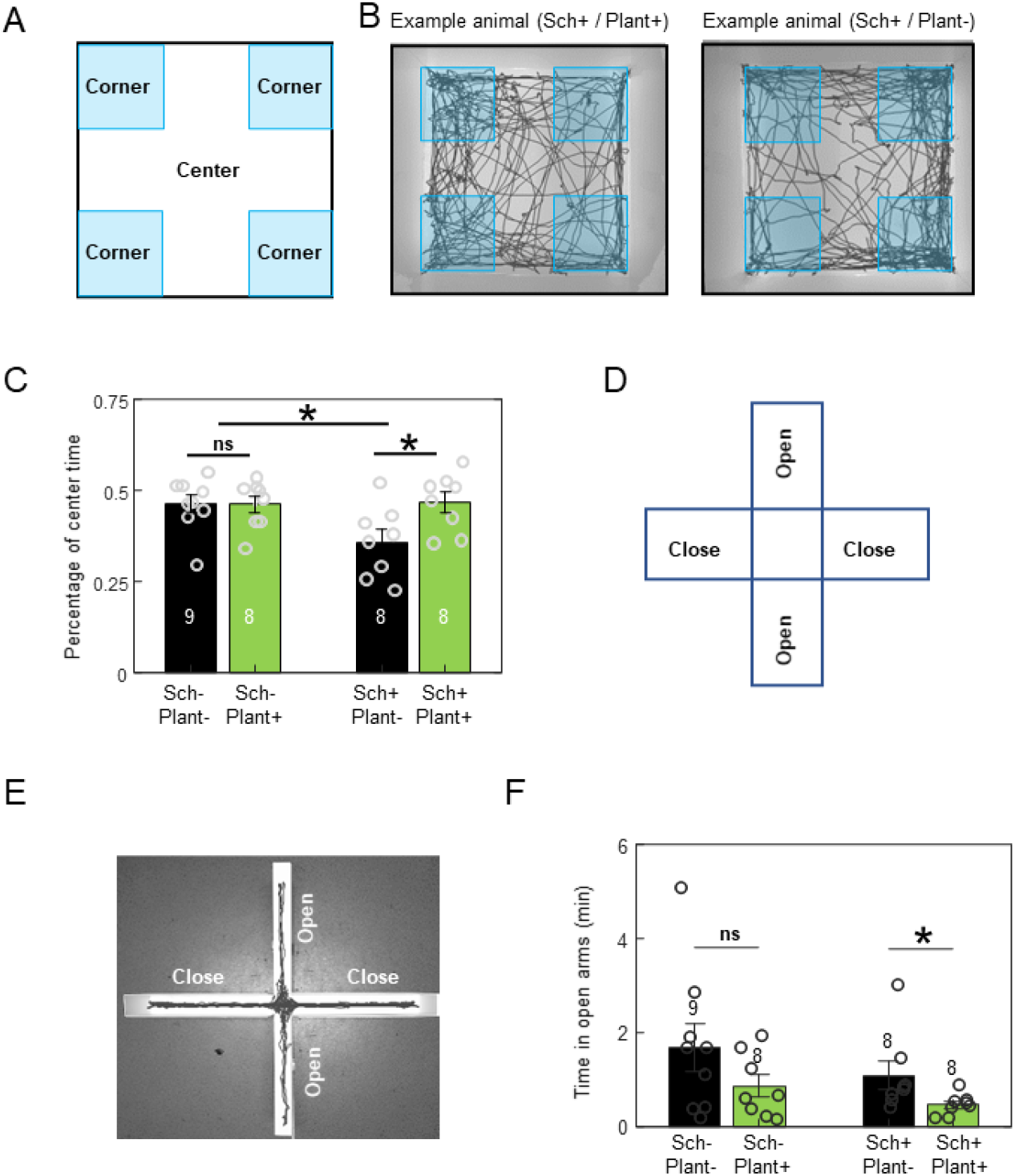
Plant exposure decreases the anxiety of schizophrenic mice in open field. (A) A diagram showing the corner and center segments of the open field which were used to define the anxious behavior. (B) The 8-minute walking paths of two representative mice in the open field test. The cyan-shaded areas label the corner segments of the open field. Left, a schizophrenic animal with plant exposure; Right, a schizophrenic animal without plant exposure. (C) The percentage of time mice from each of the 4 groups spent running in the center segment. The schizophrenic mice showed significantly less time spent in the center segment than that of WT animals, and plant exposure increased the time spent in the center. (D) A diagram showing the elevated plus maze. (E) The walking trace of a representative mouse in the elevated plus maze. (F) The time mice from each of the 4 groups spent in the open arms of the elevated plus maze.

The anxiety level of rodents was also evaluated using the elevated plus maze test, where anxious individuals usually spend more time in the closed arms than in the open arms. So, we also tested the anxiety changes of schizophrenic mice that were exposed to plants using the elevated plus maze (**Fig. 3D-3E**). Surprisingly, the anxiety level of schizophrenic mice was marginally lower compared with WT animals. In addition, the animals of both WT and schizophrenic with plant exposure spent significantly less time exploring in the open arms (**Fig. 3F**). So, plant exposure did not decrease the anxiety of schizophrenic mice in the elevated plus maze. As the mice exposed to plants were raised, we observed them frequently running or hiding within the plants. This may have led them to develop a preference for enclosed environments, resulting in the decrease of running time in the open arms which could not explained by increasing anxiety. Therefore, plant exposure reduced anxious behavior of schizophrenic mice only in the open field condition, but not in the elevated plus maze.

### Plant exposure ameliorates the social interaction deficits of schizophrenic mice

Most individuals with schizophrenia show social-related symptoms, such as social withdrawal and reduced communication, etc. Therefore, we also evaluated the social interaction of the 4 groups of mice via the three-chamber social test. The test includes 3 phases: habituation, social ability and social novelty (**Fig. 4A and 4B**). In the habituation phase, there is no mouse in either pen cup, and the test subject is allowed to freely explore the empty three-chamber box. The habituation phase is usually used to test whether the tested mouse has a preference for either of the two cups in opposite corners of the three-chamber box. In the social ability phase, one mouse is put in pen cup 1 and no mouse in pen cup 2. This phase is used to test whether the tested mouse has a stronger motivation to interact with other mice instead of the object (pen cup 2). In the social novelty phase, a new mouse is then introduced into pen cup 2. The WT usually has a stronger motivation to interact with the new mouse which is called social novelty (**Fig. 4A**).

**Figure 4.**
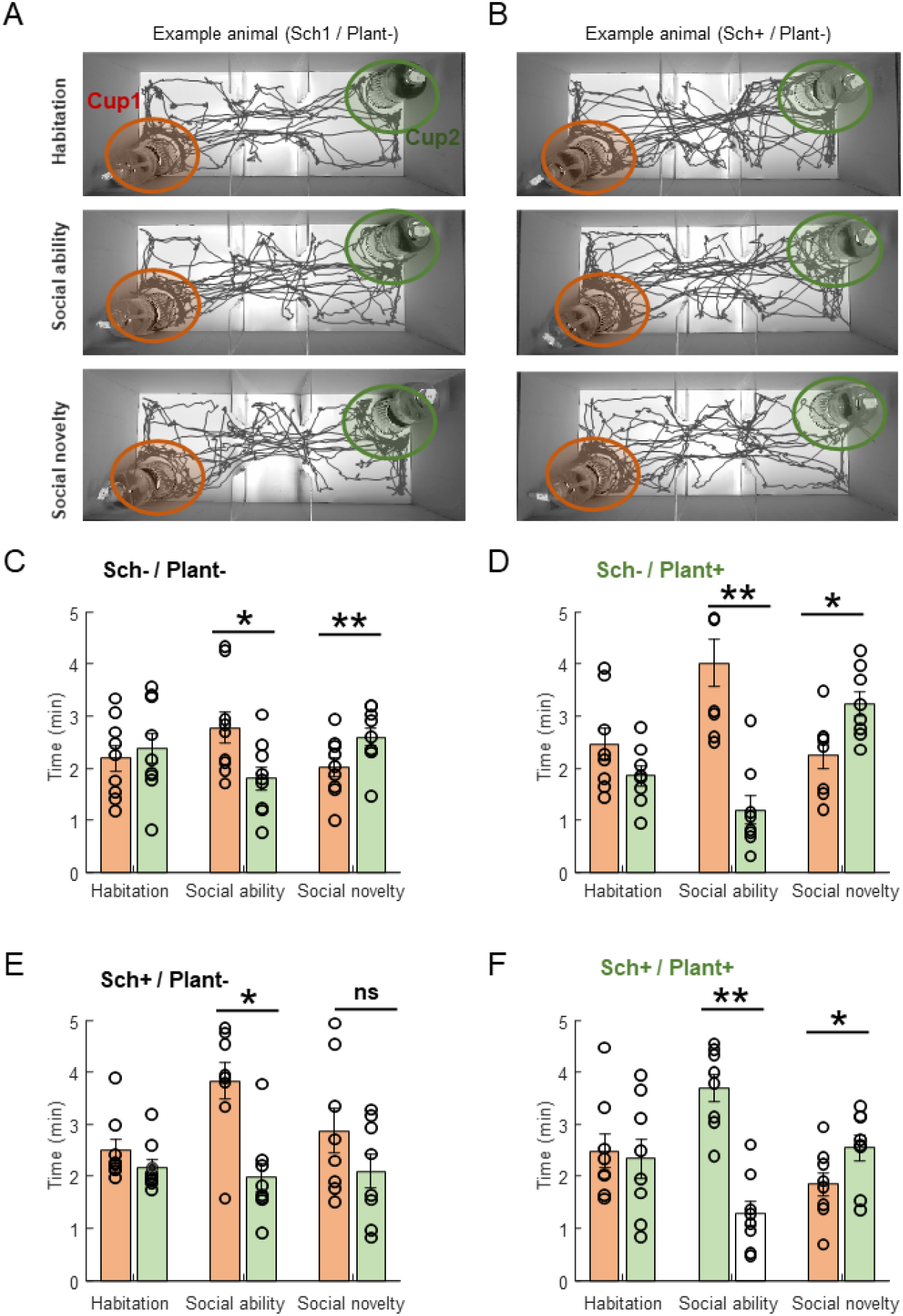
Plant exposure ameliorates the social interaction deficits of schizophrenic mice. (A) The 10 min walking trace of one example mouse (a WT animal with plant exposure) in the three-chamber box. The test includes 3 phases: habituation (Upper), social ability (Middle) and social novelty (Bottom). (B) The 10 min walking trace of 1 example mouse (a schizophrenic animal without plant exposure) in the three-chamber box. (C-F). The social behaviors of the 4 groups of animals. C, Sch-/plant-; D, Sch-/plant+; E, Sch+/plant-; F, Sch+/plant+. The interaction time of the testing mouse with the mice in the cup or the cup itself was measured by the time in the shade regions. Plant exposure improves the social novelty behaviors of schizophrenic mice.

As shown in Figures 4A and 4C, wild-type (WT) animals not exposed to plants spent similar amounts of time interacting with both cups during the habituation phase. They showed a preference for the cup containing the animal during the social ability phase and favored the cup with the novel animal during the social novelty phase (**Fig. 4A and 4C**). WT animals exposed to plants displayed similar behavior, preferring the cup with the animal during the social ability phase and the cup containing the novel animal during the social novelty phase (**Fig. 4D**). Importantly, The WT animals with plant exposure showed a larger interaction time difference between the animal and the empty cup in the social ability phase (**Fig. 4D**) in contrast to WT animals raised without plants (**Fig. 4C**). Similar to previous studies, the schizophrenic mice without plant exposure showed normal social ability, indicated by a preference for the animal rather than the empty cup in the social ability phase, while also demonstrating abnormal social behavior in the social novelty phase, indicated by similar interaction times between the novel animal and used animal (**Fig. 4E**). Interestingly, the schizophrenic mice with plant exposure showed both normal social ability and social novelty behaviors in the three-chamber social tests (**Fig. 4F**). Additionally, schizophrenic mice with plant exposure also showed a larger interaction time difference between the animal and the empty cup in the social ability phase (**Fig. 4F**) when compared to the schizophrenic mice that were not exposed to plants (**Fig. 4E**). So, plant exposure consistently improves the social interaction of mice in the social novelty phase and rescues the social novelty deficit in the schizophrenic mice.

### The c-Fos expression deficits in the hippocampus and prefrontal cortex of schizophrenic mice were partially restored by plant exposure

Previous studies have shown that abnormal prefrontal cortex^15-17^ and hippocampus^18,19^ circuits contribute to the abnormal cognitive behaviors of schizophrenia. To determine whether exposure to plants could reverse abnormalities in the prefrontal cortex and hippocampus, we performed c-Fos staining after mice from each of the 4 groups completed all three phases of the three-chamber social test. (**Fig. 5**). C-Fos is an immediately early gene which usually used as biomarker of neuronal activity. Compared the wild-type animals, schizophrenic mice showed significantly higher c-Fos expression in the prefrontal cortex (**Fig. 5A-5C**). Exposure to plants did not alter c-Fos expression in wild-type mice. However, it did slightly lower the c-Fos expression in the prefrontal cortex of schizophrenic mice, compared to the schizophrenic mice without plants (Fig. 5A-5C). Compared to WT mice, the c-Fos expression in the hippocampus of schizophrenic mice was significantly lower. (Fig. 5D-5F).

**Figure 5.**
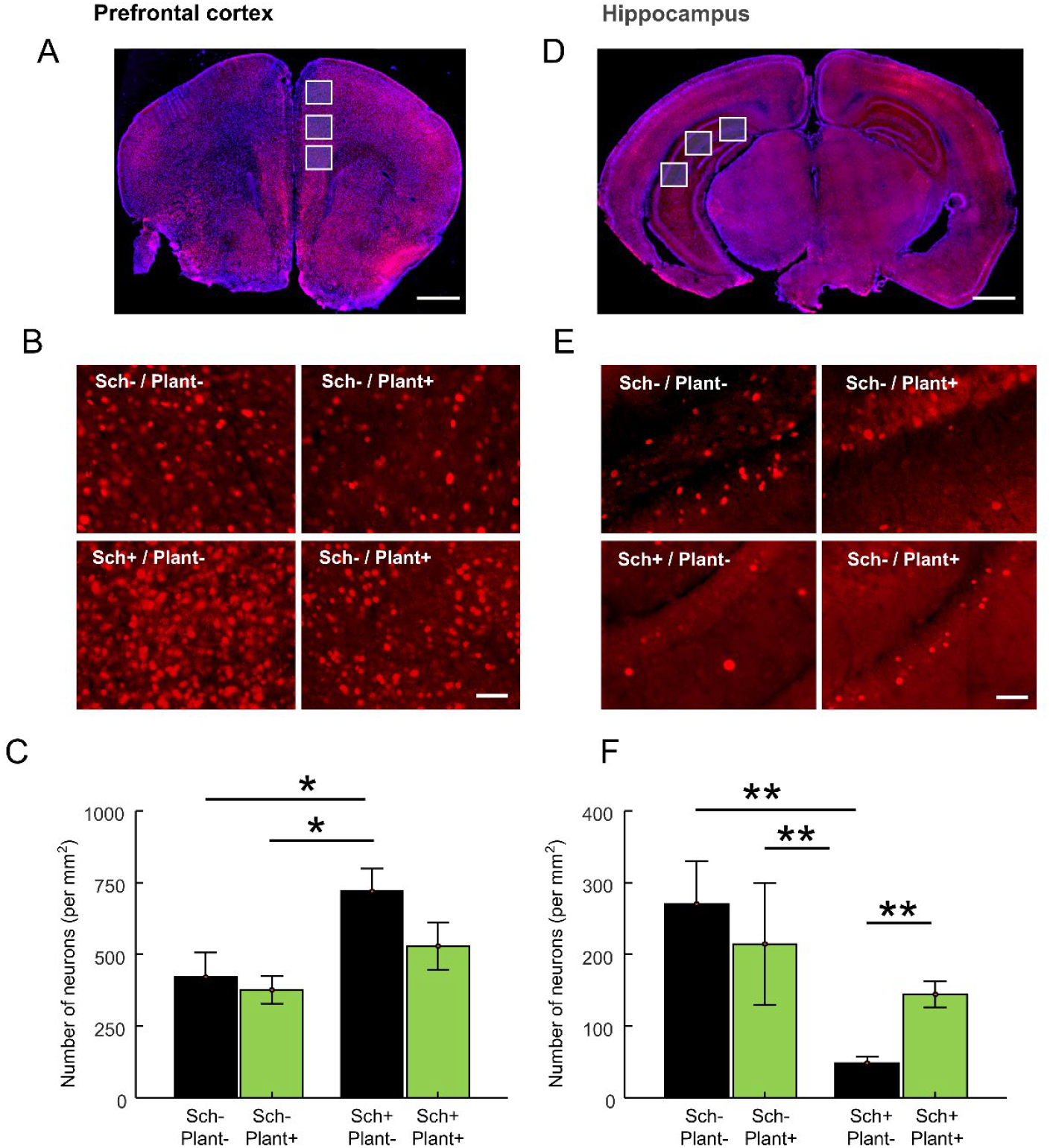
Plant exposure partially rescues c-Fos expression deficits in the hippocampus and prefrontal cortex of schizophrenic mice. (A) and (D), example images showing the overall expression of c-Fos in the whole coronal section, including the prefrontal cortex(A) and hippocampus(B) after three-chamber social tests. scale bar, 1mm. (B) and (E), example images showing the expression of c-Fos in the 4 groups of animals in the prefrontal cortex(B) and hippocampus(E). scale bar, 25 um. (C) and (F), Bar graphs display the average number of c-Fos positive cells in the prefrontal cortex(C)and hippocampus(F)across the 4 groups of animals.

More importantly, plant exposure completely rescued the c-Fos expression deficit in the hippocampus of schizophrenic mice (**Fig. 5F**). So, plant exposure can partly lower the high c-Fos expression in the prefrontal cortex and rescue the c-Fos expression deficit in the hippocampus of schizophrenic mice, which may contribute to the behavioral rescue.

## Discussion

In the present study, we first raised the mice in the environments containing two kinds of plants to induce plant exposure. Then, we investigated whether exposure to plants could mitigate negative or cognition-related deficits in individuals with schizophrenia by employing a range of behavioral tests. We found plant exposure can improve animals’ social (**Fig. 4**) and emotional-related deficits (**Fig. 3**), but do not influence the locomotive behaviors in schizophrenic mice. In addition, we tested the c-Fos expression in the hippocampus and prefrontal cortex of schizophrenic mice and found plant exposure can partly lower the high c-Fos expression in the prefrontal cortex and rescue the c-Fos expression deficit in the hippocampus of schizophrenic mice, which may contribute to behavioral rescue.

Genetic and environmental factors are two key factors in the etiology of schizophrenia ^20^. Schizophrenia is highly heritable, especially indicated by twin studies^21,22^ in which the risk of schizophrenia increased from 1% (in normal population) to 20%-45% (in twins)^22-24^. With the development of genetic tools, many risk genes have been identified including DISC1^25-27^, reelin^28-30^, Neuregulin^28^, COMT^28^, etc. It is worth noting that schizophrenia is a multigene disease, which was confirmed by a transcriptome study from the prefrontal cortex (PFC)^17^. In this study, the authors found there are 50 risk genes which organized as a network to function in neuronal migration, transcriptional regulation, transport, synaptic transmission and signaling^17^. Apart from the risk genes, abnormal environments in prenatal or early development are also one of the major factors of schizophrenia^31^. For example, schizophrenia is associated with some infections before birth^32^ where the infections affect fetal brain development^33^. Additionally, individuals living in urban environments have a higher risk of developing schizophrenia compared to those in rural areas. Recent research has highlighted that both environmental and genetic factors are crucial in the development of schizophrenia and often interact with one another.^31^. Although having different etiology, all cases of schizophrenia share similar symptoms^20,24,28^, and how schizophrenia can be treated is still a challenge, especially for negative and cognitive symptoms.

In the last 6-7 decades, many antipsychotic drugs such as SGAs, clozapine, lumateperone, risperidone, olanzapine, and quetiapine have been developed to treat schizophrenia^34,35^. Most of these drugs interact with various neurotransmitter receptors including the dopamine receptor^36,37^ and ion channels^15^. However, most of these drugs have strong side effects^24,38^ and mainly effective for treating the positive symptoms of schizophrenia^34,35^ or emotion-related symptoms^13^. Because schizophrenia is ^34,35^strong associated with negative and cognitive symptoms and poor functional outcomes for these antipsychotic drugs, effective interventions for these domains are urgently needed^24,38^.

As individuals with schizophrenia frequently experience social challenges such as withdrawal from social interactions^1,2^ several new treatment strategies have been implemented in clinical practice. These strategies include increasing community engagement, providing family psychoeducation, offering supported employment, and implementing social skills training^20^. However, many of these treatments have demonstrated only limited effectiveness. Under natural conditions, humans and animals live together with many different kinds of plants, forming symbiotic relationships. Several previous studies have suggested that exposure to green spaces can reduce the risk of schizophrenia to some extent in a dose-dependent manner^4 5-7^, and may also potentially shorten psychiatric hospitalizations^8^. Clinically, some patients showed decreased anxiety and depression after plant exposure^8,39-41^. However, all of the studies are epidemiology investigations and could not address the mechanism of plant exposure and mental disease. So, using an animal model to dissect the effect of plant exposure on schizophrenia is very important. In the present study, we made a schizophrenia mice model using long-term administration of MK-801^9,10^ and tested their social, emotional and locomotive behaviors. We found the mice displayed deficits of social and anxious behaviors after 2-3 weeks of recovery (**Fig. 3 and Fig.4**), but showed normal locomotor activity (**Fig. 2**). Importantly, social deficits observed in the three-chamber social test were eliminated following exposure to plants (**Fig. 4**). Meanwhile, the increased anxiety of schizophrenic mice in the open field condition is completely rescued with plant exposure (**Fig. 3**). However, the schizophrenic animals showed a small decrease in the time spent in the open arms compared to the control animals, and the animals with plant exposure exhibited an overall decrease in the time spent open arms compared with the animals without plant exposure (**Fig. 3**). We observed the animal’s behaviors as they were raised together with the plants, and found the animals often hid inside the plants and some mice even dug in the soil. As mice’s instinctive behavior, habits of staying in the hidden regions of their environment will become stronger during plant exposure. Thus, the reduced time spent in the open arms could not be explained by the increased anxiety. An alternative explanation is that animals raised with plants may be less anxious, but the contrast between their familiar environment and the elevated plus maze was heightened by the exposure to plants.

In this study, the addition of plants into the environment also introduced the factor of environmental enrichment, where novelty objects are present in the living space of the mice models. One explanation of the results is that environmental enrichment, not plant exposure, could be the cause of the mice’s cognitive recovery. Many studies have addressed the behavioral recovery in schizophrenic models induced by MK801 after environmental enrichment^42^. These studies proved that some cognitive impairment in rodent models could be alleviated by environmental enrichment, including learning and memory^43-45^, and sensory-motor gating^46^ in the new object recognition test, Morris water maze, Object-in-place test, and PPI test^42^. However, we did not find any reference that environmental enrichment can restore social impairments in any schizophrenic models. So, the restoration of the social and emotion-related symptoms of schizophrenic mice could not be explained by environmental enrichment. In summary, plant exposure could improve the social and emotion-related symptoms of schizophrenic mice.

Next, we tried to dissect which brain regions were involved in the behavioral restoration. As one of the most important immediate early genes, c-Fos is typically used as biomarker of neuronal activity and plasticity^47^, and is usually used to screen target brain regions in relation to special behaviors^47^. Additionally, c-Fos can be used to study the long-term neuroactivity of target brain regions, unlike the methods of transient neuroactivity recording of electrophysiology recording^48^ and calcium imaging^37,49^. Usually, the prefrontal cortex in schizophrenic individuals displays long-term high neuroactivity^37^, which is easily tested with c-Fos expression^37^. So, we examined c-Fos expression in the hippocampus^18,19^ and prefrontal cortex^15-17^ of schizophrenic mice, which are two key brain regions involved in the disease. We found that plant exposure partially reduced the elevated c-Fos expression in the prefrontal cortex and restored the c-Fos expression deficit in the hippocampus, which may contribute to the behavioral rescue (**Fig. 5**). However, transient neuroactivity recording during social behavioral tests are necessary in future to dissect transient changes during social interaction.

Although our results indicated that plant exposure can be a viable method to treat clinical symptoms of schizophrenia, there are several limitations in the present study. First, there are several types of mouse models for schizophrenia, including developmental, drug-induced, lesion-induced, and genetic manipulation models. So, it is still unknown whether plant exposure could also treat schizophrenia in other kinds of animal models. Second, there are many different kinds of plants and we only tested rosemary and *Epipremnum aureum* in the present study. Other plants including flowers, grass and xylophyta, etc. will form different plant exposure environments and should be considered in the future. Finally, because of the issue of time, deep mechanism studies, including studies of differences in transient neural activity, differences in gain and loss of function, and the effects of plant exposure duration are still ongoing. Despite these limitations, the present work provides a new method to treat the symptomatic deficits in schizophrenia and this method may be generalized to other psychotic diseases.

## Acknowledgments and Funding

This work was supported by STI2030-Major Project and 2022ZD0205000 (2022ZD0205003) and 2021ZD0204100; the Natural Science Foundation of China (32170991, 32071097 and 32371074), National Key Research and Development Program of China under Grant (2023YFB4705500),Zhejiang Province Natural Science Foundation of China (LD22H090003) and the MOE Frontier Science Center for Brain Science & Brain-Machine Integration, Zhejiang University.

## Supplemental information

Supplemental Information includes four supplemental figures.

## Data availability

Although the data files are large the original data that support the findings of this study are available upon reasonable request.

